# Compost Teas: market survey and microbial analysis

**DOI:** 10.1101/2022.08.05.503013

**Authors:** Marie Legein, Wannes Van Beeck, Babette Muyshondt, Karen Wuyts, Roeland Samson, Sarah Lebeer

**Affiliations:** Department of Bio-science Engineering, University of Antwerp, 2000, Antwerp, Belgium

**Keywords:** compost tea, microbiome, biocontrol, beneficial microbes

## Abstract

Compost teas (CTs) are fermented watery compost extracts that can be made with a wide variety of organic ingredients. These extracts are applied on plants to promote plant growth and increase protection against pathogens. Several studies have shown that the live micro-organisms present in these teas play an important role in plant protection. Still, our knowledge on the identity and functionality of bacterial communities in CTs remains limited. Moreover, CTs are a heterogeneous group, as various combinations of ingredients, fermentation parameters and application guidelines are possible and little is known on the current practices in commercially available products. Consequently, also the microbial communities in these commercially available CTs are unknown. Here, we surveyed 68 CTs on ingredients, claims and guidelines regarding preparation and application. This revealed that most products were sold in dried form which needed to be rehydrated by the customers. Moreover, kelp, humic acid, sugar and molasses, were common ingredients. Additionally, we investigated the bacterial communities of eight products, covering different ingredients and preparations, using sequencing and culture-based techniques. Bacterial communities were remarkably similar at genus level, with *Pseudomonas, Massilia, Comamonas, Chryseobacterium, Brevundimonas* and *Acinetobacter* as most prevalent genera. The genus *Pseudomonas* was more abundant in CTs that were prepared without aeration. Moreover, differences between liquid and dry products revealed the importance of formulation to secure the viability of the desired microbes. This study has given insight in common practices in CTs on the market, their microbial composition, and highlighted several discrepancies with current literature.

**Importance:** Compost teas are fermented watery compost extracts which are used in professional and amateur horticulture to promote plant growth and suppress plant diseases. Beneficial microbes present in these extracts are suggested to play an important role in the mode of action. However, little is known on the identity and functionality of the bacteria in compost teas. Moreover, compost teas are a heterogeneous group of products that can be made with numerous combinations of ingredient and fermentation parameters and little is known on current practices in products available on the market. Therefore, we surveyed these practices in 68 commercial products and for a subset of these, we explored their bacterial communities using both culture dependent and independent techniques. This study has unveiled the common practices on the market, their microbial communities, and highlighted several discrepancies between commercial practices and scientific research.

## Introduction

Crop protection and growth-promoting products are essential in ensuring a sufficient and consistent food production to a growing world population. However, current use of these products cause a range of adverse effects to human health and to the environment (United Nations Environment program 2021). Therefore, the demand for sustainable alternatives is increasing (Nishimoto 2019). One promising and fast-growing field of research is focussing on microbes as a sustainable solution to fight of pathogens and to improve plant health. Within this field, compost teas could be an underexplored source of beneficial microbes.

Compost teas (CTs) are watery fermented compost extracts that are typically applied on the rhizosphere and/or the phyllosphere to improve plant growth and/or plant protection. These extracts are assumed to be rich in microbes as well as in nutrients, such as nitrogen, phosphorous and potassium (Ingham.E.R. 2005; S. Scheuerell and Mahaffee 2002). Moreover, several studies have shown that the micro-organisms present in the CTs play a crucial role in the mode of action, since sterilization of the teas with heat or filtration usually results in a complete or partial loss of phytopathogen disease suppression (Li et al. 2020; C. C.G. St. Martin and Brathwaite 2012; S. J. Scheuerell and Mahaffee 2006; Koné et al. 2010; Dionne et al. 2012). Still, the micro-organisms present in the compost teas and responsible for the disease inhibition are not yet well characterized. In addition, the fermentation processes driving the final CTs are very diverse and poorly defined. Amongst the factors that can vary between productions are the ingredients, the fermentation time and the aeration during fermentation. This diversity in fermentation parameters results in a wide range of CT products available on the market (S. Scheuerell and Mahaffee 2002).

The aim of this study was to gain better insight into the ingredients, guidelines and claims of commercially available CTs and to identify key micro-organisms present in the final product. Therefore, a diverse set of 68 products, which were available on the market, were surveyed based on their ingredients, their preparation and application guidelines, and claims related to the products. In addition, the bacterial communities of eight CTs, covering different ingredients and preparations, were analyzed in the lab using 16S rRNA amplicon sequencing and culturing techniques.

## Materials and methods

### 1. Market survey

The online publicly available information from 68 products that were on the market and described as CTs, was assembled and analyzed. These products were found by using the key words ‘compost tea’, ‘compost thee’, ‘compost tee’, and ‘thé de compost’ in Google’s search engine on March 2020. These 68 products came from 40 different companies and nine different countries. The information analyzed included the preparation method, the form in which the products were sold, the ingredients used in the products, the application guidelines, and the claims made.

### 2. Microbial community composition of the final compost tea products by 16S rRNA amplicon sequencing

The microbial communities of eight CTs (**Error! Reference source not found.**), made from six products of which we had received a sample, were analyzed using 16S rRNA amplicon sequencing. These products were prepared according to the manufacturers guidelines. Products 1B and 4A were prepared both aerated (ACT) and non-aerated (NCT), as both preparation methods were advised. Aeration was done by bubbling air through the suspension using an aquarium air pump. NCTs were left undisturbed during fermentation.

The DNA of CTs was extracted using the PowerFecal DNA Isolation Kit (Qiagen). Each CT was extracted twice using different volumes of starting material as the microbial biomass of the teas was unknown, i.e., 100μl and 250μl CT was added to 750μl powerbead solution (PowerFecal DNA isolation kit). Next, the kit manufacturer’s instructions were followed.

The extracted DNA was subjected to barcoded PCR amplification with 25 cycles, amplifying the V4 region of the 16SrRNA gene using Phusion High-Fidelity DNA polymerase (ThermoFischer Scientific) and barcoded primers (IDT), as described by Kozich et al. (2013). The PCR amplicons were purified using AmpureXP (Agencourt) and quantified using Qubit®3.0 Fluorometer (Life Technologies). The amplicons were pooled in equimolar concentrations to obtain an amplicon library and loaded onto a 0.8% (mass/vol) agarose gel. The amplicons were cut out and purified with the NucleoSpin gel and PCR cleanup kit (Macherey-Nagel). Finally, the amplicon library concentration was measured using Qubit®3.0 Fluorometer, diluted to 2nM, and sequenced on the Illumina MiSeq platform using 2×250 cycles at the Center of Medical Genetics Antwerp (University of Antwerp).

Processing and quality control was done with the R package DADA2 version 1.4.0 (Callahan et al. 2016). In brief, reads with more than two expected errors and undetermined bases were removed. Sample inference using the DADA2 error model was performed and reads were merged. The denoised reads, i.e. amplicon sequence variants (ASVs), were then paired and chimeras removed. The ASVs were classified using the EzBioCloud 16SrRNA database (version mtp1.5, update 2018.05) (O. S. Kim et al. 2012). Finally, ASVs classified as *Eukarya* were removed. The resulting ASV tables were further analyzed in R, using an in-house package, tidyamplicons (https://github.com/SWittouck/tidyamplicons). Visualizations were made using ggplot2 (Wickham 2009).

### 3. Sample collection and preparation for culture-based microbial analyses

In parallel to the sequence-based analysis, bacteria were isolated from the same CTs (**table 1**), with two exceptions, product 1A was also prepared anaerobically and plated out (but not sequenced), and product 4A was not plated out. Freshly prepared CTs were plated out on three media: Man-Rogosa-Sharpe (MRS, Difco), reasoner’s 2 agar (R2A, Roth), and Beech-leaf-medium (BM) (Scheuerl et al. 2020). BM was made by boiling 50g dried beech leaves collected from the litter layer of a *Fagus sylvatica* stand in the Middelheim park (Antwerp, Belgium, 51°10’45.0”N 4°24’46.3”E) in 500ml of phosphate-buffered saline (PBS) during 30 minutes. The liquid was filtered to remove particles (using a paper coffee filter, approx. mesh size 20mm), agar was added (final concentration 15g/l), and the medium was autoclaved. All media were supplemented with cycloheximide (final concentration 0.1g/l) to suppress the growth of fungi. The inoculated agar plates were incubated at room temperature (ca. 22°C) during 72h. After incubation, colonies with different morphologies were picked and sub-cultured until pure cultures were obtained. These colonies were putatively classified up to genus or species level by amplifying and sequencing the 16S rRNA gene. Hereto, the 16S rRNA gene was amplified by PCR using primers 27F (5’-AGAGTTTGATCCTGGCTCAG-3’) and 1492R (5’-GGTTACCTTGTTACGACTT-3’). Amplicons were sequenced by a Sanger sequencing service (VIB Genetic Service Facility, University of Antwerp). The sequences were compared to the 16S rRNA sequence database from EZbiocloud on 03/06/2020.

**Table 1:**
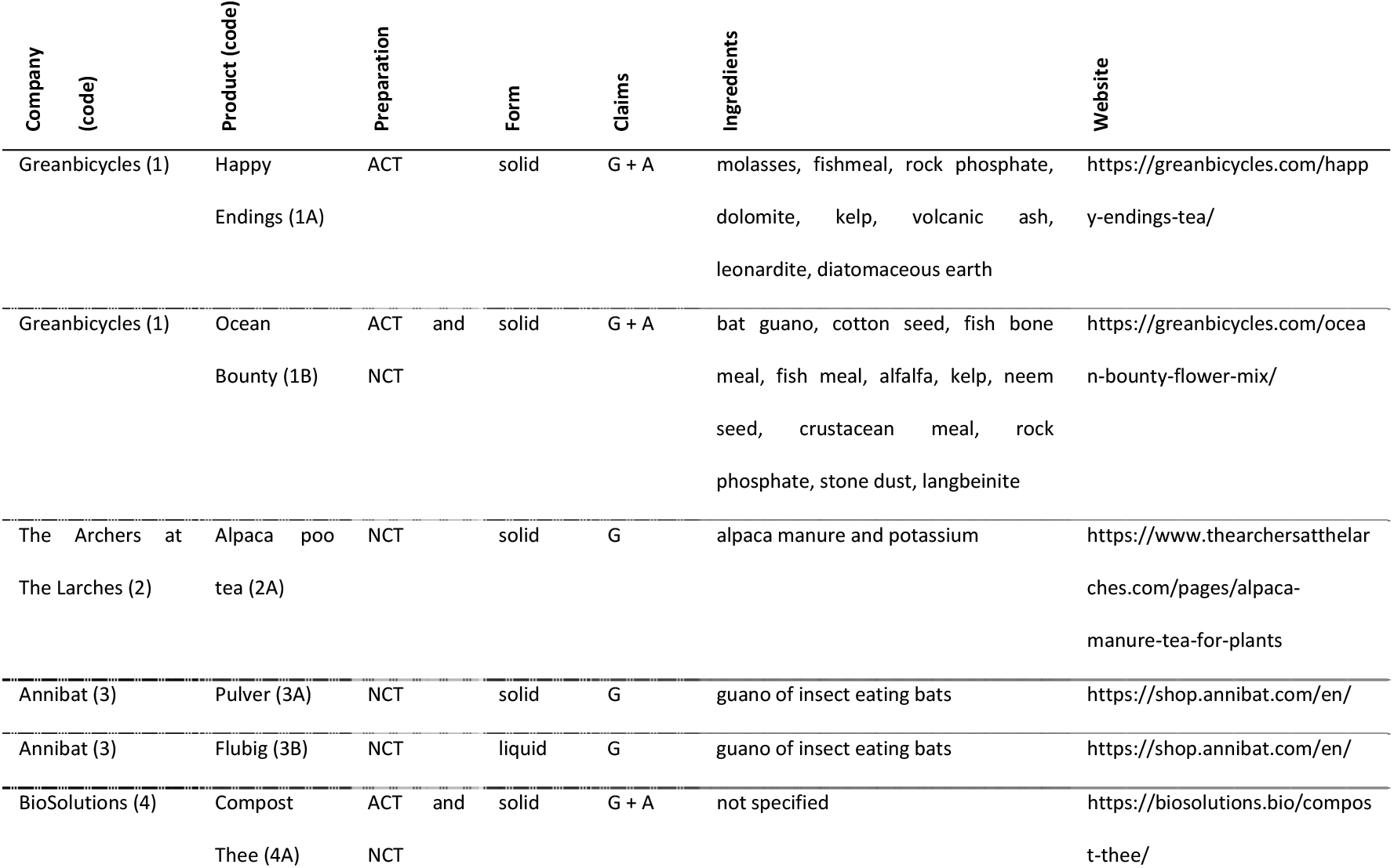
Overview of products selected for microbial analysis. All companies and products are undisclosed and thus identified by a code. Products are prepared based on the guidelines on the product (aerated (ACT) and/or non-aerated (NCT) compost tea). The table also includes information derived from the market survey: Form in which the product is sold (solid or liquid), claims (growth-promoting (G) and/or antipathogenic (A)), and ingredients in the product.

**Table 2:** 23 isolates, isolated from the compost teas and their most probable taxonomy. Identification was done using the 16S rRNA gene sequence and the EZBiocloud database. Three different media were used to isolate bacteria: Man-Rogosa-Sharpe (MRS), reasoner’s 2 agar (R2A), and beech leaf (BL) medium. Preparation methods were aerated (ACT) or non-aerated (NCT). The company and product codes correspond to the products in table 1, the similarity (% similar nucleotides) and completeness (%) of the 16S rRNA gene sequences with respect to full-length sequences in the EZBiocloud database are provided.

| Company - product code | Preparation | Top hit taxon | medium | Similarity (%) | Completeness (%) | top hit taxonomy |
| --- | --- | --- | --- | --- | --- | --- |
| 1-A | ACT | *Exiguobacterium sibiricum* | MRS | 100 | 95 | Bacteria;Firmicutes;Bacilli;Bacillales;Exiguobacterium_f;Exiguobacterium |
|  | ACT | *Glutamicibacter arilaitensis* | MRS | 100 | 95 | Bacteria;Actinobacteria;Actinomycetia;Micrococcales;Micrococcaceae;Glutamicibacter |
|  | ACT | *Glutamicibacter halophytocola* | MRS | 99 | 94 | Bacteria;Actinobacteria;Actinomycetia;Micrococcales;Micrococcaceae;Glutamicibacter |
|  | NCT | *Enterobacter hormaechei subsp. xiangfangensis* | MRS | 100 | 95 | Bacteria;Proteobacteria;Gammaproteobacteria;Enterobacterales;Enterobacteriaceae;Enterobacter;Enterobacter hormaechei |
|  | NCT | *RIBP_s (Bacillus sp.)* | MRS | 100 | 94 | Bacteria;Firmicutes;Bacilli;Bacillales;Bacillaceae;Bacillus |
|  | NCT | *Prolinoborus fasciculus* | R2A | 100 | 95 | Bacteria;Proteobacteria;Gammaproteobacteria;Pseudomonadales;Moraxellaceae;Prolinoborus |
| 1-B | ACT | *SJOH_s (acinetobacter sp.)* | BL | 100 | 93 | Bacteria;Proteobacteria;Gammaproteobacteria;Pseudomonadales;Moraxellaceae;Acinetobacter |
|  | ACT | *Exiguobacterium sibiricum* | MRS | 100 | 93 | Bacteria;Firmicutes;Bacilli;Bacillales;Exiguobacterium_f;Exiguobacterium |
|  | NCT | *Bacillus megaterium* | BL | 100 | 96 | Bacteria;Firmicutes;Bacilli;Bacillales;Bacillaceae;Bacillus |
|  | NCT | *Bacillus safensis subsp. safensis* | BL | 100 | 95 | Bacteria;Firmicutes;Bacilli;Bacillales;Bacillaceae;Bacillus;Bacillus safensis |
|  | NCT | *Carnobacterium inhibens subsp. inhibens* | MRS | 100 | 100 | Bacteria;Firmicutes;Bacilli;Lactobacillales;Carnobacteriaceae;Carnobacterium;Carnobacterium inhibens |
|  | NCT | *Carnobacterium inhibens subsp. inhibens* | MRS | 100 | 100 | Bacteria;Firmicutes;Bacilli;Lactobacillales;Carnobacteriaceae;Carnobacterium;Carnobacterium inhibens |
|  | NCT | *Enterococcus lactis* | MRS | 100 | 100 | Bacteria;Firmicutes;Bacilli;Lactobacillales;Enterococcaceae;Enterococcus |
|  | NCT | *Enterococcus lactis* | MRS | 100 | 100 | Bacteria;Firmicutes;Bacilli;Lactobacillales;Enterococcaceae;Enterococcus |
|  | NCT | *Enterobacter quasiroggenkampii* | R2A | 100 | 96 | Bacteria;Proteobacteria;Gammaproteobacteria;Enterobacterales;Enterobacteriaceae;Enterobacter |
| **2-A** | NCT | *Enterococcus faecium* | MRS | 100 | 74 | Bacteria;Firmicutes;Bacilli;Lactobacillales;Enterococcaceae;Enterococcus |
|  | NCT | *Enterococcus hermanniensis* | MRS | 100 | 95 | Bacteria;Firmicutes;Bacilli;Lactobacillales;Enterococcaceae;Enterococcus |
|  | NCT | *Enterococcus mundtii* | MRS | 100 | 95 | Bacteria;Firmicutes;Bacilli;Lactobacillales;Enterococcaceae;Enterococcus |
| **3-A** | NCT | *Glutamicibacter arilaitensis* | BL | 100 | 95 | Bacteria;Actinobacteria;Actinomycetia;Micrococcales;Micrococcaceae;Glutamicibacter |
|  | NCT | *Glutamicibacter arilaitensis* | BL | 100 | 96 | Bacteria;Actinobacteria;Actinomycetia;Micrococcales;Micrococcaceae;Glutamicibacter |
|  | NCT | *Glutamicibacter arilaitensis* | BL | 99 | 95 | Bacteria;Actinobacteria;Actinomycetia;Micrococcales;Micrococcaceae;Glutamicibacter |
|  | NCT | *Glutamicibacter mishrai* | MRS | 100 | 95 | Bacteria;Actinobacteria;Actinomycetia;Micrococcales;Micrococcaceae;Glutamicibacter |
|  | NCT | *Glutamicibacter mishrai* | MRS | 100 | 93 | Bacteria;Actinobacteria;Actinomycetia;Micrococcales;Micrococcaceae;Glutamicibacter |

## Results

### 1. Market survey

We surveyed the preparation methods, ingredients, and claims of 68 CTs, which were available online on the market on March 2020. A first observation was that, although CTs should be applied on the plant in liquid form, the majority of CTs in our survey (83% or 57/68) were sold in dry form. These dry products needed to be prepared by the consumer by adding water, with or without aeration. Most products sold in solid form (n=57) needed to be prepared without aerating the mixture (67% or 38/57). For the remaining 33% (19/57) of the solid products, it was advised to aerate, using for example an air pump. For four products (1A, 1B, 4A and 32A), both aerobic and anaerobic preparation methods were advised. Regarding the products sold in liquid form, half of these mentioned that they had been prepared without aeration. For the other half, the preparation method was not described.

For almost all products studied here, it was advised to apply them on the rhizosphere, only for two products it was advised to apply only on the phyllosphere. For little over half of the samples (54% (or 37/68)), both rhizosphere and phyllosphere application was suggested. The majority (78% or 53/68) of products had claims related to growth-promoting effects, 21% (14/68) of products had claims related to both growth-promoting and antipathogenic effects, and for one product only antipathogenic effects were claimed.

The ingredients mentioned on the labels were diverse. On average, a CT contained five ingredients, the maximum was 13 and minimum one. The most frequently used ingredient was kelp: 59% (or 40/68) of products contained this ingredient (**figure 1**). The second most frequently used ingredient was non-composted plant material (56% or 38/68). Most often, these plant sources were alfalfa (21%, 14/68) and nettle (16%, 11/68). Besides these, a wide diversity of plant materials were listed in only one or two products, including oak and willow bark, valerian, dandelion, chamomile, yarrow, rapeseed, neem seed, comfrey, and soybean meal or soy hydrolysate. The third class of ingredients most often added were sources of humic acid (41%, 28/68).

**Figure 1.**
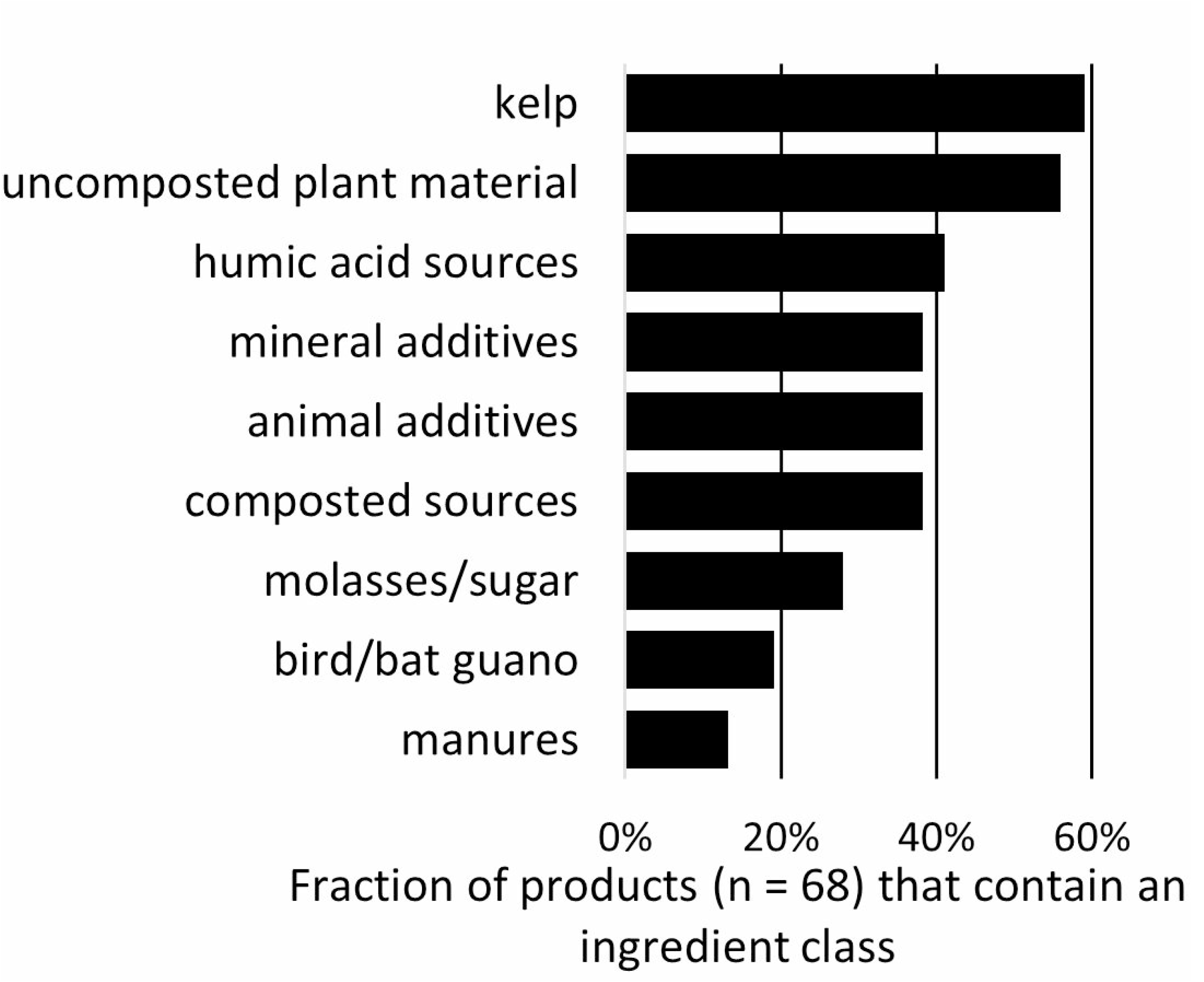
Share of products that contain the specified ingredient class. Ingredients were divided in nine classes: kelp, uncomposted plant material, humic acid sources, mineral additives, animal additives, composted sources (vermicompost and bokashi), molasses/sugar, bird/bat guano, manures.

Next, 38% (26/68) of the products contained a mineral source, including rock phosphate, rock dust, volcanic ashes, diatomaceous earth, dolomite and glauconite. In addition, various ingredients of animal origin were used as source of minerals. These include feathers, fish bones, fish meal and bone meal (manures are discussed as a separate ingredient class). About one third (32%, 22/68) of the investigated products contained mineral-rich animal-based ingredients.

Furthermore, 38% (25/68) of investigated CTs contained a source of compost. Most often, vermicompost (the product of earthworm decomposition of organic matter) was added. In two products, bokashi was used (the product of an anaerobic fermentation of organic matter, often amended with a source of microbes such as dairy products or commercially available effective microbes (Quiroz and Céspedes 2019)).

Molasses or sugars were added to 28% of investigated CTs. Finally, one fifth (19%) of CTs included bird or bat guano and 13% of products included manures from other sources, most often coming from cows, one CT reported horse manure and another alpaca manure.

### 2. Community diversity and composition of compost teas

The bacterial community composition of eight CTs, made from six products of which two were prepared both aerated and non-aerated (1B and 4A, Error! Reference source not found.), were further analyzed by amplifying and sequencing the 16S rRNA genes. Two of these products contained bat guano as sole ingredient and came from the same company, one of these was sold in liquid form (3B), and the other in solid form (3A). Both these products were prepared without aeration. Two other products (1A and 1B), also coming from the same company, contained various ingredients, such as kelp, molasses, and mineral additives. Both were prepared with aeration and additionally, as suggested on the packaging, product 1B was prepared also without aeration. A third product (4A) did not specify the ingredients and was also prepared both with and without aeration. The last product (2A) contained alpaca manure and potassium as ingredients and was prepared without aeration.

Sequencing of these samples resulted in 795,808 reads, after data processing and quality control. One sample (3B, 100μl extraction) was excluded from further analysis, as it did not contain any reads. On average, each sample contained on average 41,396 reads, and minimum 21,062 reads. The number of taxa detected in each sample ranged between 129 and 318, excluding product 3B which contained only 15 taxa, **Table 1**. The Inversed-Simpson index varied between 16 and 46, excluding again product 3B which had a much lower diversity. The diversity indices were similar for both aerated and non-aerated teas (**figure 2A**).

**Figure 2:**
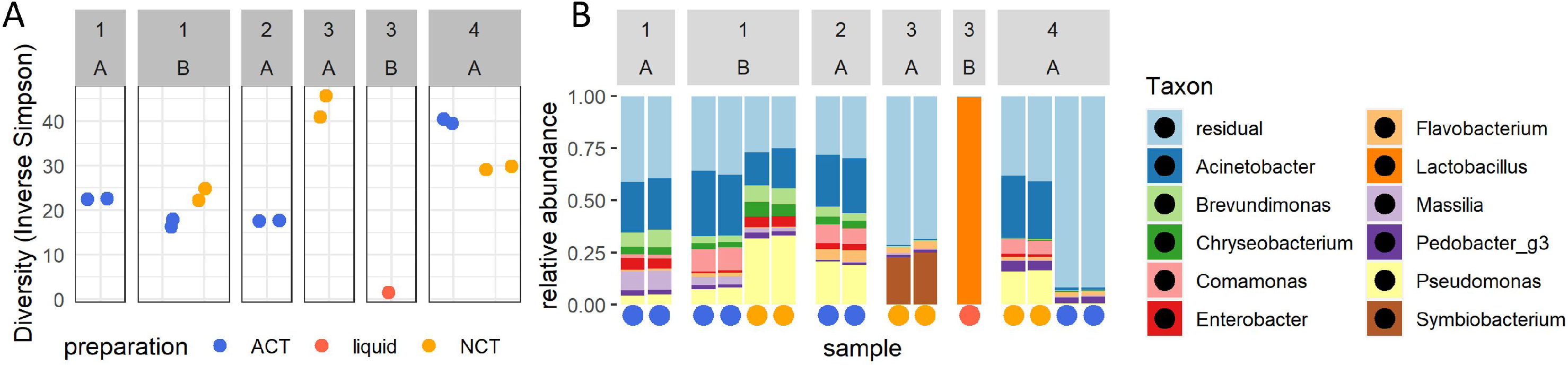
Community diversity (A) and composition (B) of eight compost teas analyzed with 16S rRNA amplicon sequencing. (A) inverse Simpson diversity indices. (B) Barplots showing the 11 most abundant genera. Each sample was extracted and sequenced twice. Samples are ordered per company and product (numbers 1-4 above, table 1), and based on preparation method, aerated (ACT, blue), non-aerated (NCT, orange), and samples sold in liquid form (liquid, red)

In our analysis, product 3B was an outlier, the only product in our analysis which was sold in liquid form, was an outlier as it had a lower diversity compared to other samples (**figure 2A**). Moreover, its community was dominated by a single taxon, classified as *Liquorilactobacillus* (**figure 2B** and **figure 3**). In contrast, product 3A, which came from the same company and was also made with bat guano as sole ingredient but sold in solid form, contained a diverse microbial community and no taxa classified as *Liquorilactobacilli* (**figure 2B** and **figure 3**).

**Figure 3:**
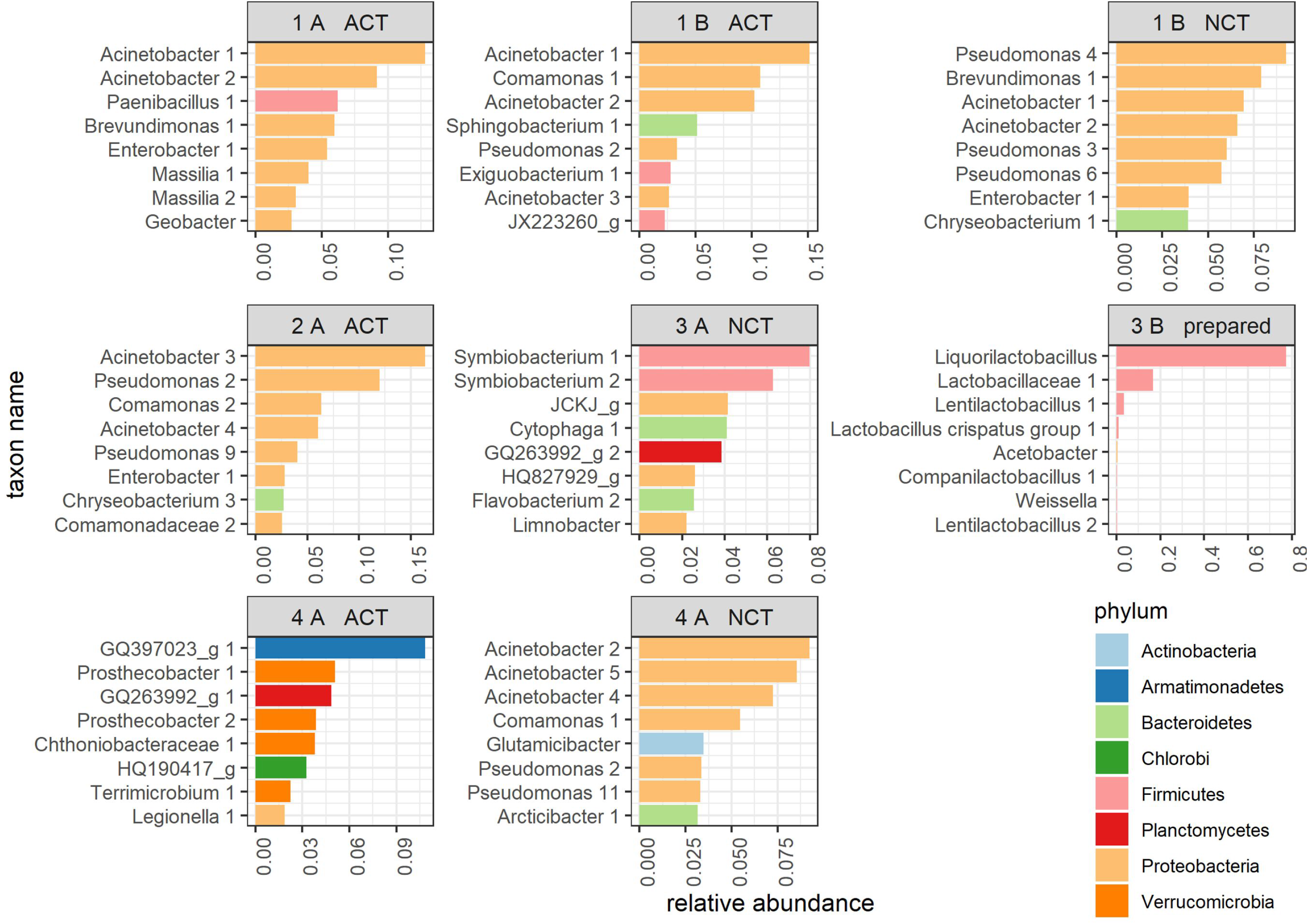
Bar charts showing the relative abundances of the eight most abundant amplicon sequence variants (ASVs) in each compost tea sample. Sample identification is indicated above each chart; company 1-4, product A-B (product list in table 1), preparation method (aerated (ACT), non-aerated (NCT) or sold in liquid form (liquid)). The bars are colored according to the phylum the ASV belongs to.

At genus level, the different CTs, excluding samples from company 3, were remarkably similar (**figure 2B**). The most prevalent genera occurring in the samples were *Pseudomonas, Massilia, Comamonas, Chryseobacterium, Brevundimonas* and *Acinetobacter.* However, at the level of amplicon sequence variants (ASVs), which roughly corresponds to species level, the different products contained different ASVs within these genera (**figure 3**). At phylum level (**figure 3**), communities were characterized by a high prevalence of *Proteobacteria, Firmicutes* and *Bacteroidetes.*

### 3. Identification of culturable bacteria in compost teas

Next to the sequence based identification of the microbial communities in CTs, also the active and culturable members of the community were identified. Twenty-three bacteria, belonging to eight different genera were cultured from different CTs and identified by sequencing the full-length 16S rRNA gene (Error! Reference source not found.).

Five isolates were classified as lactic acid bacteria and belonged to the genera *Carnobacterium* and *Enterococcus.* These lactic acid bacteria were isolated from two products, both prepared without aeration. Based on the microbial community analysis, product 3B was dominated by one ASV from the genus *Liquorilactobacillus* (**figure 3**). However, it was not possible to isolate any bacteria from this product on neither of the three used media (MRS, BL or R2A).

Bacteria from the genus *Glutamicibacter* were isolated from non-aerated product 3A and aerated 1A. Bacteria from the genus *Bacillus* were isolated from non-aerated 1A and 1B, and *Exiguobacterium* strains were isolated from the aerated version of these products. Finally, bacteria from the genus *Enterobacter,* as well as *Acinetobacter* and *Prolinoborus* (closely related to *Acinetobacter Iwoffii)* were isolated from 1B and 1A.

## Discussion

Our market analysis revealed that CTs on the market are a highly diverse group, made with a wide diversity of ingredients and fermentation parameters and that these practices not always correspond with findings from scientific literature.

Firstly, products on the market are either sold in liquid or dry form, while scientific research focusses on CTs made with fresh ingredients. The dry products need to be rehydrated and fermented by the end-users and the liquid products often only need dilution upon use. However, the microbes present on the dry starting materials are most likely not the same as the microbes on fresh material. Especially for microberich materials, such as vermicompost and other composts, drying and storage must have an unwanted effect on the microbial composition of the end product. Also long-term storage of liquid products will have a negative effect on the viability of the microbes. The differences in microbial community between similar products in dry and liquid form will be discussed later.

Next discrepancy was that one third of products (33%) needed to be prepared with aeration, while several disadvantages of aeration have been pointed out in literature. Ingram and Millner (2007) pointed out the risk of human pathogen growth in aerated conditions, as they found that populations of *E. coli, Salmonella enteritidis* and *Enterococcus* were higher in ACTs compared to NCTs. Additionally, the advantages of aeration are scarce as most studies that directly compared aerobic and non-aerobic preparation methods did not find a difference in disease control (S. J. Scheuerell and Mahaffee 2006; Al-Dahmani et al. 2003). Still, research is not conclusive on this topic, for example Seddigh and Kiani (2018) observed a better suppression of powdery mildew in a greenhouse setting with ACTs compared to NCTs and NCTs can be more phytotoxic due to the presence of organic acids (Carballo et al. 2009; Kannangara, Forge, and Dang 2006).

On the topic of ingredients, we found a high diversity of ingredients used in commercially available CTs. One remarkable observation was that the majority of products contained amendments, i.e. ingredients additional to the bulk organic material, such as kelp, humic acid, sugar, molasses, and various sources of minerals. Previous research suggests that amendments generally result in higher microbial population counts (Naidu et al. 2010; Ingham.E.R. 2005). However, this could also result in enhanced growth of unwanted bacteria such as *E. coli, S. enteritidis, Enterococcus* and fecal coliforms (Kannangara, Forge, and Dang 2006; Ingram and Millner 2007). In addition to the risk of unwanted bacteria, Scheuerell and Mahaffee (2006) did not observe a significant difference in disease suppression of ACTs amended with or without various amendments, including molasses, kelp, trace minerals, humic acids and rock dust. In conclusion, the common presence of amendments in commercial CTs calls for an assessment of the risk of human pathogens in CTs on the market.

Further popular ingredients were vermicompost, manure and guano. It has been suggested that CTs produced with vermicompost result in higher disease suppression, possibly due to a higher bacterial diversity in vermicomposts compared to other composts (Ingham.E.R. 2005) as the vermicomposting process is non-thermophilic and earthworm-associated microbes are added (Pathma and Sakthivel 2012). However, this advantage might be irrelevant for commercial applications as these microbes would lose their viability upon storage, both in dry as in liquid form.

Regarding manures, several studies have demonstrated positive antipathogenic and growth-promoting effects of CTs prepared with different manures (Litterick et al. 2004; S. J. Scheuerell and Mahaffee 2006; McQuilken, Whipps, and Lynch 1994; Evans, Palmer, and Metcalf 2013; Koné et al. 2010). Although, literature points out the risk of *E. coli* in swine manure (Kannangara, Forge, and Dang 2006). In our assessment of the market, this hazard was not present as none of the products contained swine manure.

Next, we analyzed the microbial composition with culture dependent and independent techniques, as the microbes within these extracts are assumed to play an important role in the efficacy of CTs. A subset of eight commercially available CTs were analyzed and these products contained various ingredients and fermentation parameters. The bacterial communities in these products were remarkably similar at phylum and genus level, but more distinct at ASV level. Although Ingham (2005) hypothesized that ACTs are generally more diverse, no studies could be found comparing the diversity between aerobic and non-aerobic preparation methods.

Here, we did not observe a consistent difference in alpha-diversity between aerated and non-aerated CTs and our results were similar to previous studies on NCTs (Mengesha, Gill, et al. 2017; Mengesha, Powell, et al. 2017).

Amongst the most abundant taxa were several ASVs belonging to the *Pseudomonas* genus. Interestingly, in products 1B and 4A which were prepared both as ACT and NCT, the relative abundance of *Pseudomonas* was higher in the NCT compared to the ACT samples. Thus non-aerated conditions might favor the growth of *Pseudomonas* in compost teas. The benefits of the presence of *Pseudomonas* is uncertain as several strains are known biocontrol agents with inhibitory activity against a range of phytopathogens (e.g. *Pseudomonas chlororaphis* MA342 and *Pseudomonas* sp. DSMZ13134), while other strains are pathogenic for a range of plants [e.g. *P. syringae* (Xin, Kvitko, and He 2018)]. Despite the presence in the sequence data, no viable *Pseudomonas* were isolated from these teas. Therefore, further research is needed to determine the viability and biocontrol activity of *Pseudomonas* in NCTs.

Next to the sequence-based approach, we also explored the active and culturable bacteria in these CTs. We specifically focused on the presence of lactic acid bacteria as several studies have described the presence of these bacteria in CTs (Naidu et al. 2010; Mohd Din et al. 2018; M. J. Kim et al. 2015) and their potential beneficial role on the plant (Lamont et al. 2017). However, to our knowledge, there are no studies that further identified and characterized viable lactic acid bacteria in CTs. Here, we isolated lactic acid bacteria from the genera *Carnobacterium* and *Enterococcus. Carnobacteria* have been frequently isolated from various environments, in particular cold environments, as well as living fish and food derived from animal products (Leisner et al. 2007). The isolates from the genus *Enterococcus* isolated from 1B and 2A cause some concern, as *Enterococci* are generally described as opportunistic human pathogens.

Remarkably, no viable bacteria could be isolated from product 3B, a liquid product from bat guano. However, based on the sequence-based approach, this product was dominated by one lactic acid bacterium belonging to the genus *Liquorilactobacillus.* These bacteria normally grow well on the used MRS medium (Zheng et al. 2020), thus the bacteria in this product probably no longer viable or below the detection limit (10 CFU/ml). It is known that viable cell counts decrease rapidly in prepared CTs after a few months (Naidu et al. 2010) and this illustrates again the issue to keep the desired microbes alive in commercial CTs. Amending CTs with humic acid and yeast extract has been reported to improve the viability of microbes in liquid CTs but these ingredients were not present in this product. In contrast to the liquid product 3B, the solid product 3A from the same manufacturer, which was also made with bat guano as sole ingredient and without aeration, did not contain *Lactobacilli* in high abundances but was composed of a more diverse community instead. Thus, bacteria in the liquid product lost viability, while fermentation of the dry solid product did not result in the same lactic acid dominated end-product. While it is unknown which bacterial composition was most desired in these two CTs, the difference between the dry and liquid product shows the importance of formulation to preserve the desired micro-organisms. One possible solution to preserve desired microbes is the addition of a dried microbial starter to dried products (Broeckx et al. 2016). To our knowledge, none of the surveyed CTs added dried beneficial microbes to their products, nor did we find any research articles investigating this approach.

Next to these lactic acid bacteria, bacteria from the genus *Glutamicibacter* were isolated from three products that were prepared both with and without aeration. Members from this genus are often isolated from soil, roots and leaves and have been shown to promote plant growth under saline conditions (Qin et al. 2018), to have antagonistic properties against *Erwinia tracheiphila* (Fu, Olawole, and Beattie 2021), and improve the degradation of night soil compost (human manure) (Borker et al. 2020). These *Glutamicibacter* strains are thus of interest for further research on plant growth-promoting and/or biocontrol functions.

Next, also several isolates from the genus *Bacillus* were isolated from NCTs and could be important members of the CT community. Bacilli form spores to persist in adverse environments (Kovács 2019), which is expected to be an advantage for persisting in dry CTs. Moreover, several *Bacilli* are known for plant growth-promoting and biocontrol properties (Hashem, Tabassum, and Fathi Abd_Allah 2019), including strains that have been isolated from CTs (On et al. 2015). Additionally, bacteria from the genus *Exiguobacterium,* also members of the *Bacillaceae* family, were isolated from two ACTs. This genus does not have the advantage of producing spores, but has been described to have plant growth-promoting properties and antagonistic effects against various phytopathogens (Kasana and Pandey 2017).

Finally, bacteria from the genus *Enterobacter,* as well as *Acinetobacter* and *Prolinoborus* (closely related to *Acinetobacter lwoffii)* were isolated from 1B and 1A. These genera frequently occur in compost and CTs (S. Scheuerell and Mahaffee 2002; Chaney C. G. St. Martin et al. 2020), and were also abundant in the amplicon sequencing data. However, some members of these genera are known to cause disease in humans. Therefore, further research is needed to assess the risk of human pathogens.

In conclusion, this study has unveiled the common practices for CTs available on the market and highlighted several discrepancies between practices in commercial CTs and scientific research. Importantly, some of these call for an assessment of the risk of human pathogens in commercial CTs. Moreover, we pointed out the effect of storage and drying on the microbial communities in the end product and suggest that to further explore formulation to preserve wanted microbes. Moreover, we isolated and identified the live and culturable bacteria present in these teas. These strains could play a role in the efficacy of compost teas as a whole and could also be valuable to develop as individual biocontrol or biostimulant strains.

## Acknowledgements

The authors would like to thank Lukas Six, Dries Van Hoof, and Charline Klünder for collecting the CT products and setting up the experiments in the context of their bachelor thesis project. Furthermore, we would like to thank the CT producers for their cooperation and contributions of samples.

## Data availability statement

The sequencing data were deposited in the European Nucleotide Archive (ENA) under study accession number PRJEB43218 (https://www.ebi.ac.uk/ena/browser/view/PRJEB43218).

